# Controlling Drug Partitioning in Individual Protein Condensates through Laser-Induced Microscale Phase Transitions

**DOI:** 10.1101/2024.03.12.584573

**Authors:** Axel Leppert, Jianhui Feng, Vaida Railaite, Tomas Bohn Pessatti, Cecilia Mörman, Hannah Os-terholz, Filipe R.N.C. Maia, Markus B. Linder, Anna Rising, Michael Landreh

## Abstract

Gelation of protein condensates formed by liquid-liquid phase separation (LLPS) occurs in a wide range of biological contexts, from the assembly of biomaterials to the formation of fibrillar aggregates and is therefore of interest for biomedical applications. Soluble-to-gel (sol-gel) transitions are controlled through macroscopic processes such as changes in temperature or buffer composition, resulting in bulk conversion of liquid droplets into microgels within minutes to hours. Using microscopy and mass spectrometry, we show that condensates of an engineered mini-spidroin (NT2repCT^YF^) undergo a spontaneous sol-gel transition resulting in the loss of exchange of proteins between the soluble and the condensed phase. We find that liquid spidroin condensates absorb visible light, which enables us to control sol-gel transitions of individual droplets through laser pulses. Fluorescence microscopy reveals that laser-induced gelation significantly alters the interactions between droplet proteins and small molecules, which allows us to load single droplets with an anticancer drug. In summary, our findings demonstrate direct control of phase transitions in individual condensates opening new avenues for functional and structural characterization.

**SYNOPSIS TOC:** The liquid-to-solid transitions of phase-separated protein condensates are challenging to control. Leppert *et al*. show that condensates of engineered mini-spidroins gelate at slightly elevated temperatures. Using high-energy laser pulses at wavelengths that are absorbed by the droplets, the authors induce sol-gel transitions in single droplets. These gelated droplets are chemically stable and exhibit an increased ability to sequester drug molecules.

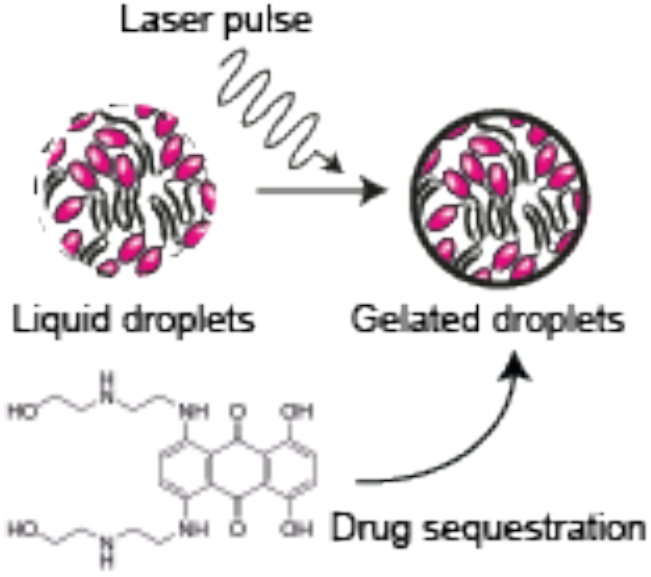

## INTRODUCTION

Liquid-liquid phase separation (LLPS) is a wide-spread phenomenon in nature, connecting physics, chemistry, biology, and material science.^1,2^ The regulated assembly of proteins through LLPS is an important mechanism for the formation of biologically active condensates within living cells, but also the basis for solid biomolecular structures like squid beak and spider silk, shifting the physical state of the protein assembly.^3–5^ During LLPS, proteins separate from a uniform single phase into a protein-rich, condensed phase and a protein-scarce, dilute phase. Depending on molecular interactions and external conditions, the material properties of protein droplets dynamically change and range from liquid-like to dynamically arrested gel-or glass-like.^6,7^ However, many proteins that form biomolecular condensates can undergo multiple phase transitions where the liquid-like properties are lost and the droplets exhibit gel-like features, such as the inability to fuse, increased chemical stability, and non-spherical shapes.^8^ In several human diseases, proteins that have a strong propensity for β-sheet formation, like Fused in Sarcoma, tau, and α-Synuclein, can be assembled into liquid droplets, which can protect them from aggregation. ^9,10^ In some instances, these droplets undergo sol-gel transitions as an intermediate step in the conversion from liquid droplets to fibrils.^11–13^ In other instances, phase transitions can be of functional importance. By adding phase separation-promoting sequence tags, Wei *et al*. were able to assemble functional organelles with colocalized enzymes in bacteria. ^14^ During spider silk spinning, protein molecules called spidroins are assembled via LLPS, which in turn prepares them to be converted into a solid fiber.^4,15–18^ Spider silk-inspired peptides have furthermore been used to generate phase-separated particles. ^19,20^ Such controlled phase transitions are of importance in biomedical engineering. For example, gelation of protein coacervates can be used to generate microgels for drug delivery.^21^

The importance of phase transitions in disease and bioengineering has led to concerted efforts to control sol-gel transitions and the properties of microgels in molecular detail.^22,23^ However, phase transitions are usually triggered on a large number of droplets by exposing them to an external stimulus, like changes in temperature, ionic strength, pH, or cofactors. ^8^ Control of individual droplet transitions are achieved in microfluidics-based set-ups (for example, see ^24–26^). As a result, spatio-temporal control of droplet gelation remains challenging. However, one way to locally steer phase transitions in proteins is the use of focused light, as exemplified by the use of cryptochrome domains to engineer proteins that exhibit light-induced LLPS.^27^ Here, we demonstrate that laser pulses can be used to control sol-gel transitions in mini-spidroins that undergo functional phase transitions as part of the spinning process. By targeting single spidroin droplets with laser pulses at micrometer resolution, we are able to accelerate the gelation process in single droplets from hours to seconds and monitor the phase transition using structurally sensitive fluorescent probes. We show that gelation locally enhances the affinity for small molecules that preferentially interact with aromatic residues. Our findings open the possibility to control sol-gel transitions at the microscale to steer the recruitment of small molecule cargo.

## RESULTS AND DISCUSSION

### Fluorescence Spectroscopy and Mass Spectrometry show NT2RepCT^YF^ Microgel formation

As a test case for functional, controllable phase transitions, we turned to the designed mini-spidroin NT2RepCT, which features a conserved three-domain architecture with nonrepetitive N-terminal (NTD) and C-terminal (CTD) domains encapsulating a central region with two alanine- and glycine-rich repeat sequences.^28,29^ Importantly, NT2RepCT, as well as the isolated NTD, can be converted into β-sheet-rich hydrogels through incubation at elevated temperatures and high concentrations.^30^ Spidroin hydrogels have been used for cell culture and the release of therapeutic biologicals.^30,31^ Macroscopic liquid-to-solid transitions of NT2RepCT can be triggered by pH, shear force, as well as changes in salt concentrations or temperature, making it a suitable system to study sol-gel transitions at the microscale.^28^ We selected the NT2RepCT^YF^ variant, where all Tyr residues in the repetitive region are exchanged to Phe, which exhibits robust LLPS without affecting its ability to be spun into fibers.^32,33^ These observations can be explained by a preference for spherical assembly due to increased contributions from π-stacking. ^34^

Combining the conditions for droplet formation and gelation,^16,31,32^ we incubated NT2RepCT^YF^ at a concentration of 25 μM in 0.5 M KPO_4_, pH 8, overnight. We observed a moderate increase in droplet size, in line with previous studies,^32^ but no other morphological changes compared to fresh droplets. However, upon addition of 10% 1,6-hexanediol, a potent LLPS disruptor,^35^ fresh droplets readily dissolved, leaving some amorphous aggregates, whereas the incubated droplets remained unaffected, indicating gelation (Figure 1 a). Since spun NT2RepCT silk is rich in β-sheet structures,^36^ we tested whether this also is the case for gelated NT2RepCT^YF^ droplets. Indeed, incubation of fresh droplets with the dye Thioflavin T (ThT) yielded an increase in fluorescence over 24 h (Figure 1 b). Fluorescence microscopy showed weak ThT staining after 5 mins, indicating that the dye is recruited into the droplets, as well as strong uniform staining after 24 h (Figure 1 c). Since ThT fluorescence is sensitive to changes in viscosity, we additionally probed the conformation of NT2RepCT^YF^ in droplets using the dye pFTAA, which exhibits a specific, characteristic two-maxima emission spectrum when bound to β-sheet structures.^37,38^ We found that gelated droplets exhibit a two-maxima spectrum comparable to that of the model amyloid Aβ_42_ (Figure 1 d). Our fluorescence data suggest that gelation of NT2RepCT^YF^ droplets likely involves an increase in β-sheet content, although other structural changes cannot be excluded.

**Figure 1.**
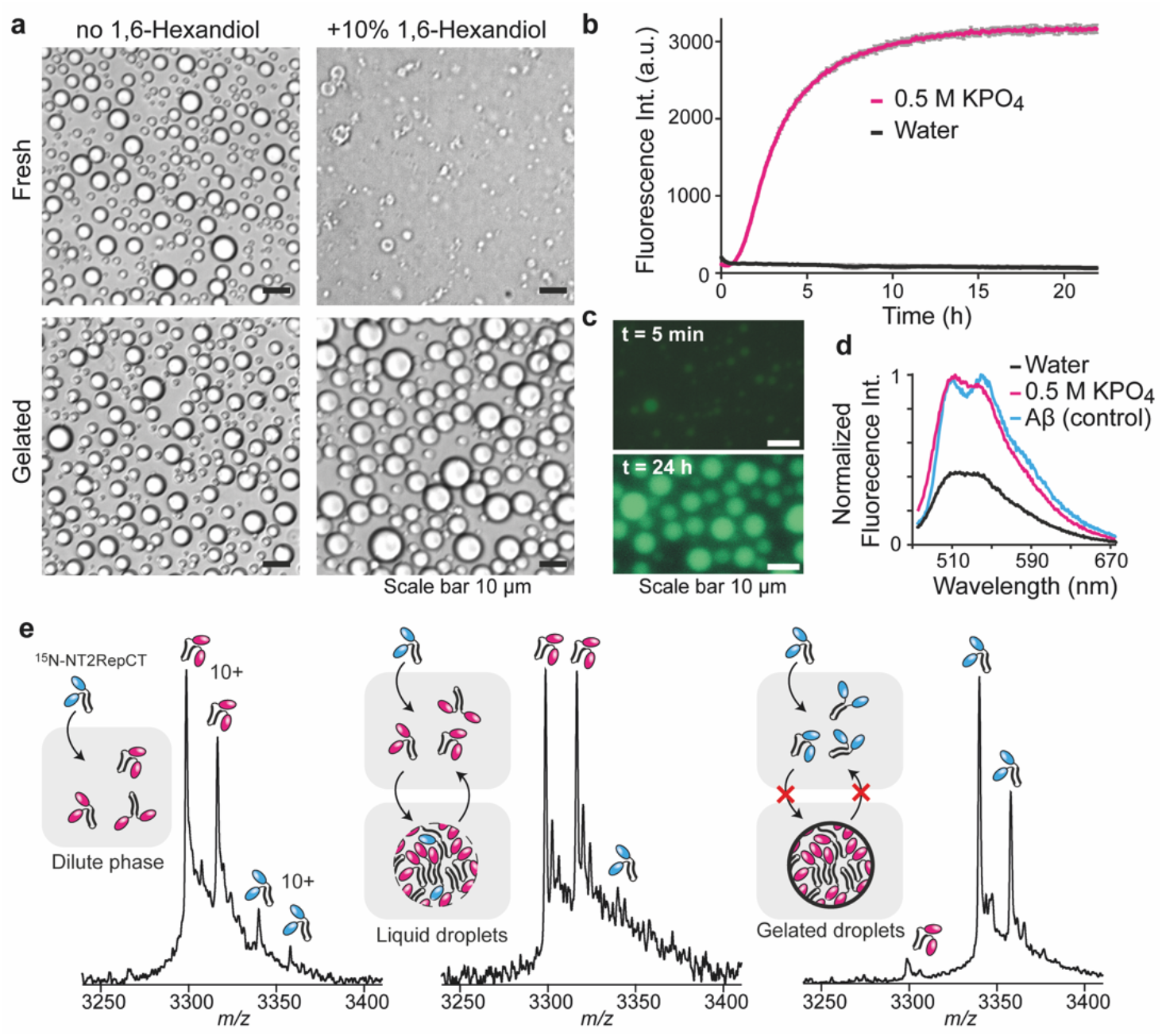
Gelation of NT2RepCT^YF^ droplets. (a) Brightfield images show that freshly formed NT2RepCT^YF^ droplets dissolve upon addition of 1,6-hexanediol (top row) but become 1,6-hexanediol-resistant upon gelation (bottom row). Scale bars are 10 μm. (b) NT2RepCT^YF^ droplets exhibit an increase in ThT fluorescence during incubation under LLPS conditions (0.5 M KPO_4_, pH 8). Data is presented as mean ± standard deviation (grey) of 4 replicates. (c) Fluorescence microscopy shows weak ThT-staining of the droplets after 5 min, as well as strong ThT-staining at the endpoint of gelation. Scale bars are 10 μm. (d) The emission spectrum of NT2RepCT^YF^ droplets after incubation (pink curve) stained with pFTAA shows the characteristic maxima indicating β-sheet formation. The spectra of NT2RepCT^YF^ in water, and of Aβ_42_ fibrils (positive control) are shown in black and blue, respectively. (e) An nMS assay shows the sol-gel transition of NT2RepCT^YF^ droplets. Left: Mass spectrum of 100 μM NT2RepCT^YF^ with 10 μM ^15^N-labeled NT2RepCT^YF^ under non-droplet conditions (100 mM ammonium acetate). Middle: Mass spectrum of ^15^N-NT2RepCT^YF^ added to fresh NT2RepCT^YF^ droplets shows rapid equilibration of the labeled and unlabeled protein in the dilute phase. Right: Mass spectrum of ^15^N-NT2RepCT^YF^ added to gelated NT2RepCT^YF^ droplets shows that the labeled protein remains in the dilute phase. The double peaks stem from formyl-methionine cleavage in the *E. coli* expression host.

Together, the resistance to 1,6-hexanediol and increase in β-sheet-specific dye fluorescence indicate the formation of macroscopic spidroin assemblies during gelation. To test this possibility, we developed a native mass spectrometry (nMS) assay based on the assumption that proteins undergo constant exchange between the condensed and the dilute phase in liquid, but not in gelated droplets. Briefly, we added ^15^N-labeled NT2RepCT^YF^ to unlabeled NT2RepCT^YF^ and determined the ratio of both proteins in the dilute phase (the supernatant) with nMS. ^15^N-labeled protein was added to 100 μM unlabeled protein at a concentration of 10 μM, below the threshold for droplet formation (Figure S1), resulting in a 10:1 ratio of unlabeled to labeled NT2RepCT^YF^ in the total protein population (Figure 1 e, left). When ^15^N-NT2RepCT^YF^ was added to freshly formed unlabeled droplets, a similar ratio of unlabeled to labeled protein was detected in the dilute phase, indicating that the labeled protein rapidly equilibrated with the unlabeled protein across the condensed and dilute phases (Figure 1 e, middle). Upon addition of ^15^N-NT2RepCT^YF^ to unlabeled gelated droplets, virtually only labeled protein was detected in the dilute phase, suggesting that it is excluded from the condensed phase (Figure 1 e, right). We conclude that incubation of NT2RepCT^YF^ under LLPS conditions converts liquid droplets to spherical microgels, which is likely accompanied by an overall increase in β-sheet content.

### Laser Pulses Induce Sol-Gel Transitions of Individual NT2RepCT^YF^ Droplets

The most-studied phase transitions occur in neurodegeneration-associated proteins such as FUS, tau, and α-Synuclein. Here, amyloidogenic segments in unstructured protein regions inside the dynamic phase-separated assemblies make stochastic contacts with each other, eventually nucleating fibrillar structures over time.^39,40^ In NT2RepCT, on the other hand, the most amyloidogenic regions are concealed in the folded terminal domains which readily change their structures following external cues.^41,42^ To better understand how the metastability of spidroins affects gelation, we turned to fluorescence recovery after photo-bleaching (FRAP) coupled with microscopy, an established tool to assess the dynamics of protein condensates.^43,44^ As first step, we sought to clarify the motility of ThT in fresh droplets. To our surprise, we found that a 20 s pulse with a laser wavelength of 405 nm at 25% laser power caused an immediate increase in ThT fluorescence intensity, which overshot the pre-bleaching fluorescence by 1.3-fold. When laser power was raised to 100%, the overshoot increased to approximately 1.7-fold with a narrow standard deviation (Figure S2). Microscopy revealed that the ThT fluorescence increase starts in the bleached spot and spreads out within 6 minutes (Figure 2 a). To test whether this behavior is specific for ThT, we repeated the experiment using the viscosity-sensitive DroProbe reagent which does not absorb at 405 nm,^45^ and additionally selected a rectangle as the laser-exposed region to test whether the shape would be retained. As with ThT, DroProbe rapidly diffused back into the bleached area, which remained rectangular for >2 minutes and exhibited a 1.5-fold over-shoot in fluorescence post-bleach (Figure 2 b). The same feature was observed for droplets composed of NT2Rep with no CTD (Figure S2). For comparison, we performed FRAP at maximum laser power with full-length tau (htau), an established amyloidogenic phase-separating protein.^11^ We observed complete DroProbe bleaching and no fluorescence overshoot (Figure S2). We then asked whether the protein itself also remained mobile after the laser pulse. We therefore assembled fresh droplets containing 0.25 μM Atto655-labeled NT2RepCT^YF^ and performed FRAP a laser wavelength of 405 nm at maximum laser power. The Atto655 fluorescence in the bleached region of the droplet did not recover, although unbleached droplets continued to fuse droplets (Figure S2). Taken together, these data suggest that the spidroins in the bleached area convert into a more viscous phase, indicating gelation. To confirm this interpretation, we repeated the experiments with already gelated droplets that were prepared in absence of any light source during incubation at 25°C for 72 h. ThT and DroProbe exhibited stronger fluorescence than in fresh droplets from the start but did not show a fluorescence overshoot after slowly diffusing back into the bleached regions (Figure 2 d and e). As expected, Atto655-labeled NT2RepCT^YF^ did not diffuse back into the bleached spot (Figure 2 f), strongly suggesting that gelation indeed abolishes the laser-induced fluorescence overshoot.

**Figure 2.**
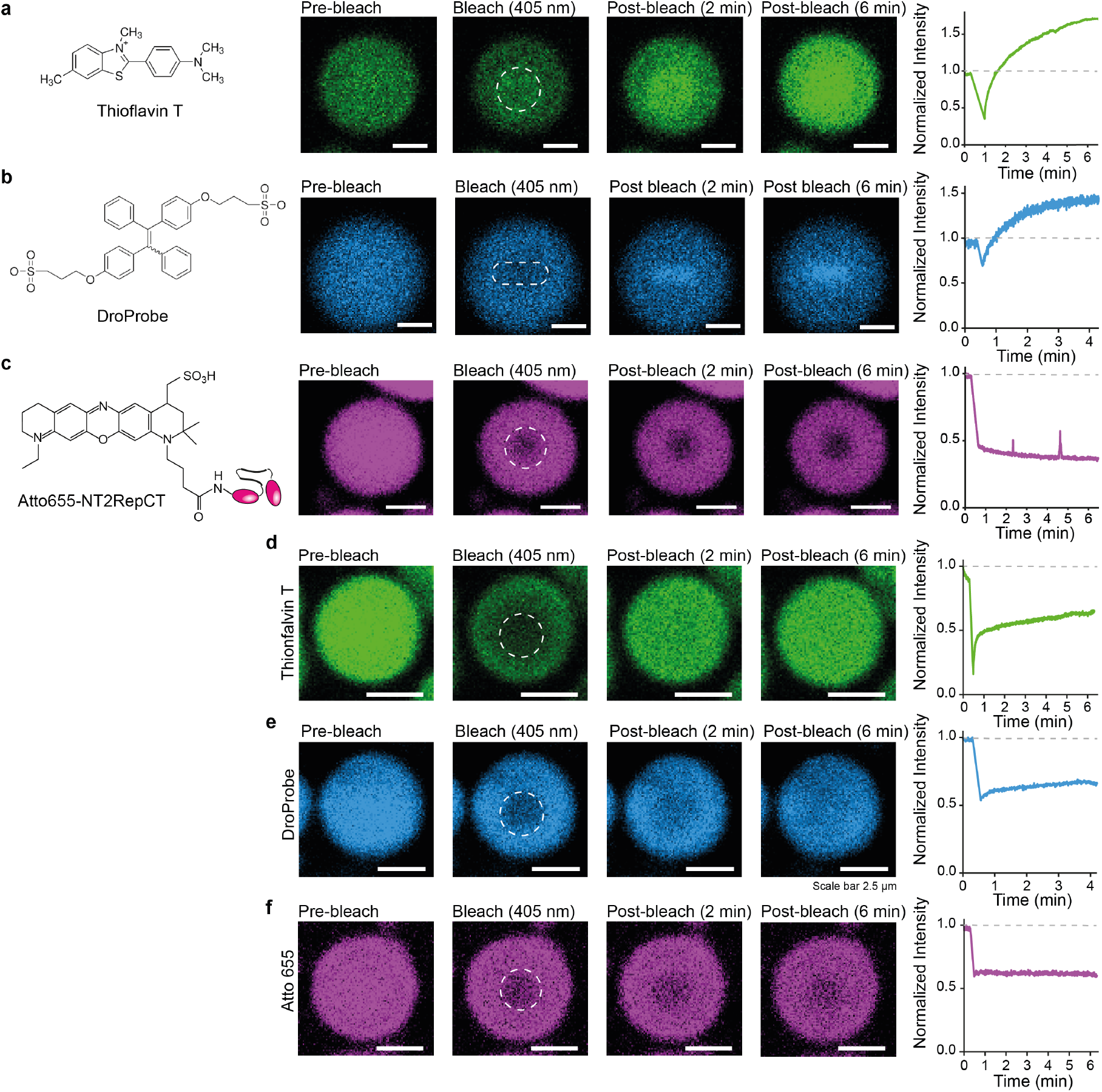
Laser-induced sol-gel transitions in individual fresh, but not gelated, NT2RepCTYF droplets. (a) The dye ThT shows an increase in fluorescence after FRAP, which spreads from the bleached area (dashed circle) to the whole droplet. (b) The viscosity dye DroProbe also exhibits increased fluorescence after FRAP, while the bleached area initially retains its rectangular shape. (c) FRAP of NT2RepCTYF droplets containing 0.25 μM Atto655-labeled protein shows no recovery in the bleached area (dashed circle). (d-f) Gelated droplets show only partial recovery of ThT (d) and DroProbe fluorescence (e), and no recovery of Atto655 fluorescence after photobleaching (f). Time-dependent fluorescence intensity plots for the center of the photobleached area are shown to the right of each series. Note that each plot refers to the droplet shown to the right. See Figure S2 for representative errors. Scale bars are 5 μm in (a,b,c) and 2.5 μm in (d,e,f).

The FRAP data suggest that laser pulses can greatly accelerate gelation of fresh spidroin droplets, reducing the time from hours to seconds. This finding appears surprising, as NT2RepCT^YF^ does not contain any chromophores that can interact with light in the visible range. Interestingly, amyloid-like fibrils absorb light at wavelengths between λ 360 - 700 nm and exhibit red-shifted fluorescence in the visible and near-IR region.^46–48^ The origin of the phenomenon, which is particularly prominent in silk fibers,^49^ is not clear, and multiple explanations have been put forward, including quantum confinement effects, electron delocalization, and charge transport.^50,51^ LLPS of spidroins promotes self-assembly and β-sheet fibrillation, although the β-sheet content may differ depending on the spidroin sequences used.^4,16,18,32^ We therefore speculated that spidroin droplets may interact with visible light in a similar manner as silk fibers. Indeed, we found that excitation at λ 405 nm and λ 561 nm (blue and green light, respectively) resulted in red-shifted fluorescence of fresh droplets, which became more intense upon gelation (Figure S3). htau droplets, on the other hand, exhibited pronounced fluorescence only at λ 405 nm excitation (Figure S3), similar to htau fibrils.^52^ The data indicate that some protein condensates, like amyloid-like fibers, may have protein-specific fluorescence properties. Importantly, the red-shifted fluorescence confirms that spidroin droplets can absorb visible light, and that some of the energy is dissipated in other ways than photon emission. It is therefore likely that the absorption of high-energy laser pulses of the same wavelength may help to overcome the energy barrier for gelation, for example through heating, since a temperature increase of 4 - 7 K is enough to trigger the assembly of NTDs into hydrogels.^30^ Our findings thus open the possibility of using spidroin domains to engineer protein condensates with laser-inducible sol-gel transitions.

### Laser-Induced Sol-Gel Transitions Enhance Drug Partitioning into Droplets

Protein condensates can sequester antineoplastic drugs by engaging their aromatic moieties in non-specific π-π and π-cation interactions.^53^ Importantly, sequestration can reduce the efficacy of these compounds by preventing them from reaching their therapeutic targets, but how such interactions are affected by phase transitions within the condensate is not known. Inducing gelation at the microscale raises the possibility to investigate how molecular interactions between drugs and individual droplets are affected by sol-gel transitions. We selected Mitoxantrone, an anthracenedione used to treat acute myeloid leukemia, prostate cancer, and multiple sclerosis. Recruitment of Mitoxantrone, which partitions into nuclear FIB1, NPM1, and Med1 condensates *in vitro*,^53^ into spidroin condensates was confirmed by monitoring its intrinsic fluorescence in fresh NT2RepCT^YF^ droplets (Figure 3 a). We then induced gelation by photobleaching with a 20 s maximum energy laser pulse and 639 nm wavelength on an individual droplet. Strikingly, we detected a 1.5-fold overshoot in fluorescence already at the end of the pulse, which subsequently increased to 2-fold within 6 minutes. No change in Mitoxantrone fluorescence was detected in the neighboring, non-bleached droplets (Figure 3 a). Unlike ThT and Droprobe, Mitoxantrone fluorescence, which stems from the rigid anthracene moiety, is independent of the chemical environment. Therefore, the increase likely indicates that the gelated droplet has a higher affinity for the drug than the surrounding liquid droplets. To confirm this interpretation, we performed a double-photobleaching experiment. A fresh, once-bleached droplet was allowed to recover before being bleached a second time. We then measured the resulting change in Mitoxantrone fluorescence. While the fluorescence recovery after the second bleach was rapid, the droplet fluorescence did not increase over the original level, indicating that the sol-gel transition induced by the first photobleaching event drives Mitoxantrone recruitment. Again, neighboring droplets exhibited no change in fluorescence (Figure 3 b). To confirm that we are indeed able to manipulate the sol-gel transition of an individual droplet, we tested the droplet stability after addition of 1,6-hexanediol. Applying our bleaching approach, we first enhanced Mitoxantrone partitioning in a single droplet (Figure 3 c). We then directly added 10% 1,6-hexanediol to the supernatant, which disrupted the integrity of all condensates, except the previously bleached droplet. Similarly, previously bleached droplets were found to be resistant to 10% formic acid (Figure S4). The preferential Mitoxantrone partitioning into the gelated droplets appears surprising. nMS reveals only very weak interactions between NT and Mitoxantrone, suggesting that binding to the folded domain is not responsible for recruitment into droplets (Figure S4). However, it was recently reported that tyrosine residues in the repeat regions of spidroins experience a shift in chemical environment during phase transitions.^54^ Furthermore, interactions with aromatic residues are responsible for drug partitioning into condensates.^53^ Although modelling structural states of the proteins in gelated droplets is not possible at this stage, we speculate that the aromatic residues in NT2RepCT^YF^ may arrange in a way that increases their affinity for aromatic compounds, possibly by enabling π-stacking interactions with the DNA intercalator Mitoxantrone.

**Figure 3.**
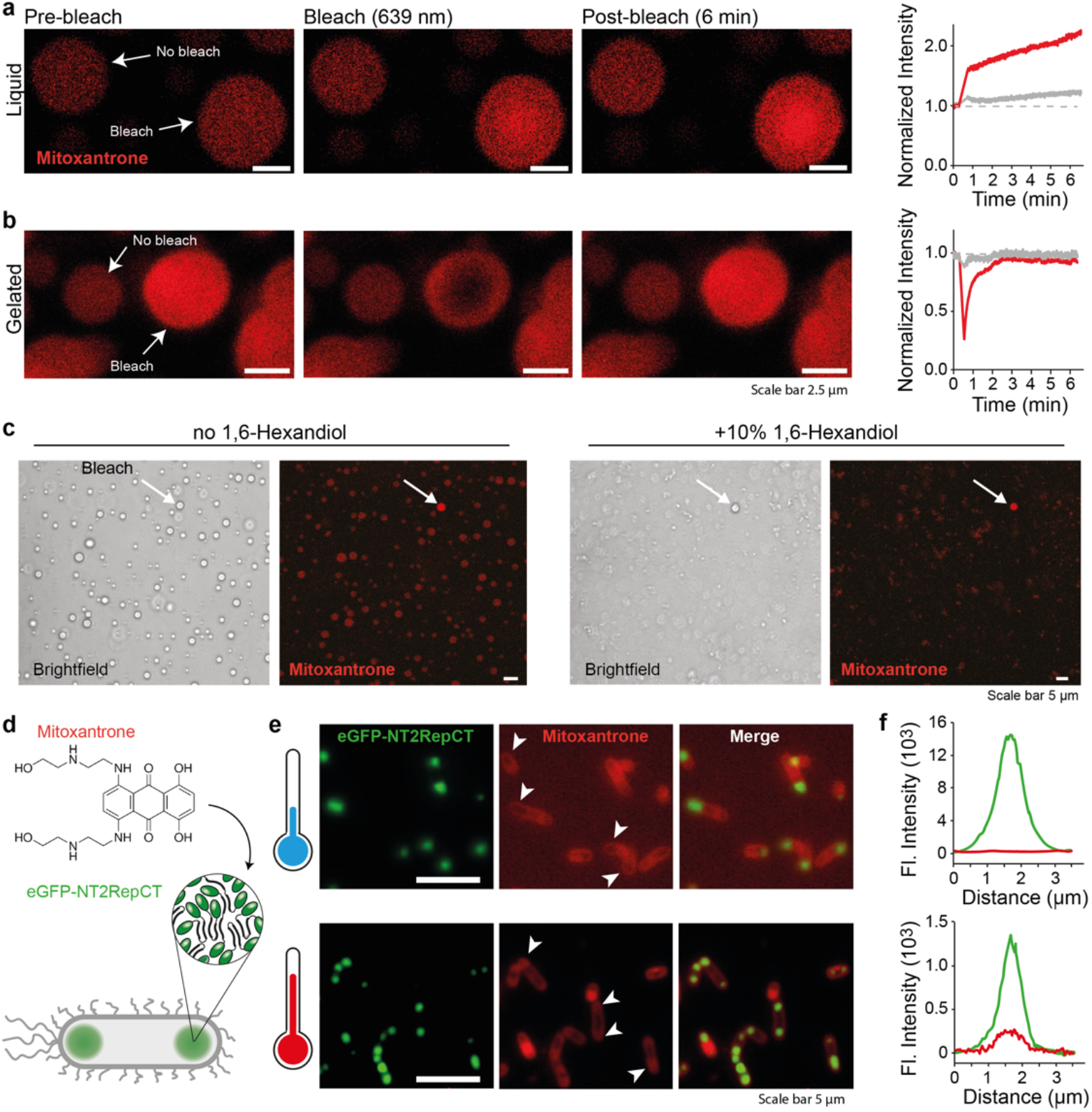
Sol-gel transitions increase drug partitioning in spidroin condensates *in vitro* and in cells. (a) Left: Fluorescence microscopy images of NT2RepCT^YF^ droplets in the presence of 25 μM Mitoxantrone (red) show low-level fluorescence, indicating recruitment into droplets. Photobleaching of a single droplet induces a strong increase in Mitoxantrone fluorescence. Right: The fluorescence intensity plot of a bleached droplet (red trace) and a neighboring, unbleached droplet (grey trace) show that Mitoxantrone fluorescence increases locally already during bleaching. (b) Photobleaching a droplet for a second time (left) does not elicit another fluorescence intensity overshoot, but shows normal fluorescence recovery (fluorescence intensity plot, right). Scale bars are 2.5 μm. (c) Photobleaching of an individual droplet (arrow) induces resistance to 1,6-hexanediol. Brightfield and fluorescence microscopy before (left) and after (right) addition of 10% 1,6-hexanediol are shown. Scale bars are 5 μm. (d) Experimental strategy for Mitoxantrone recruitment into intracellular NT2RepCT condensates in *E. coli*. (e) Top row: eGFP-NT2RepCT condensates formed during expression at 18°C show no co-localization of spidroins (green) and Mitoxantrone (red). Instead, Mitoxantrone is excluded from the condensates (white arrows) Bottom row: Expression at 37°C results in Mitoxantrone concentration at the termini of the bacteria (white arrows). Scale bars are 10 μm. (f) Fluorescence profiles of individual intracellular condensates show co-localization of Mitoxantrone and eGFP-NT2RepCT after 37°C expression (bottom), but not at 18°C (top).

Lastly, we asked whether the increase in drug partitioning upon phase transition also occurs in a cellular environment. For this purpose, we employed eGFP-tagged NT2RepCT, which forms condensates at the poles in *E. coli*.^55^ NT2RepCT droplets display the same increase in Mitoxantrone fluorescence upon laser-induced gelation as NT2RepCT^YF^ (Figure S4). Exposing the high laser energies required to induce gelation caused lysis of the cells (Figure S4). However, we previously observed that expression at 18°C, but not at 37°C, yields soluble NT2RepCT which can be purified. FRAP analysis confirmed that spidroin condensates formed at low and high temperatures are liquid and gel-like, respectively (Figure S4). To compare their ability to sequester aromatic compounds in cells, we expressed NT2RepCT at 18°C or 37°C in the presence of Mitoxantrone and assessed the colocalization of spidroin and drug using fluorescence microscopy (Figure 3 d). At low expression temperature, colocalization could not be detected, suggesting that the drug is excluded from the spidroin condensates. At high expression temperature, on the other hand, we observed eGFP-NT2RepCT assemblies which contained Mitoxantrone (Figure 3 e and f). Taken together, the data show that intracellular spidroin condensates formed at 37°C recapitulate a key feature of the laser-induced sol-gel transition.

## SUMMARY

In this study, we have demonstrated that the designed minispidroin NT2RepCT^YF^ readily undergoes a sol-gel transition following LLPS, which can be accelerated dramatically using laser pulses to allow gelation of individual droplets. We use this approach to show that the resulting microgels have an increased affinity for the antineoplastic compound Mitoxantrone both *in vitro* and in live bacteria. Micromanipulation of protein condensates is a useful tool to study their structure and function in the cellular context.^27^ The fact that mini-spidroin droplets can be gelated by laser pulses at the microscale is likely related to the ability of spidroins to convert to a fibrillar form. The fluorescence data may indicate an increase in β-sheet formation. Confirming this hypothesis requires structural characterization with single-droplet resolution, which is an emerging area in condensate research.^56^ While the use of laser pulses to induce gelation is not directly applicable to other LLPS assemblies like tau droplets, it provides a path to other light-controlled sol-gel transitions, for example through photoactivatable domains that expose fibril-forming sequences when exposed to light of a specific wavelength. With such tools, it will be possible to study the molecular effects of sol-gel transitions, both functional and pathogenic, in the cellular environment.

## ASSOCIATED CONTENT

### Supporting Information

The supporting information contains the experimental methods and Supporting Figures 1-4 (pdf).

## AUTHOR INFORMATION

### Author Contributions

AL, AR, and ML designed the study with input from JF and MBL. AL and CM produced protein. AL, VR, and OH performed microscopy and mass spectrometry experiments. JF performed live-cell imaging. TBP and FRNCM provided additional expertise. AL, AR, and ML wrote the paper with input from all authors.

## ABBREVIATIONS

LLPS, Liquid-liquid phase separation; nMS, native mass spectrometry; ThT, thioflavin T; FRAP, fluorescence recovery after photobleaching; htau, human tau protein.

## REFERENCES

(1) Shin, Y.; Brangwynne, C. P. Liquid Phase Condensation in Cell Physiology and Disease. Science (1979) 2017, 357, eaaf4382. 10.1126/science.aaf4382.

(2) Fuxreiter, M.; Vendruscolo, M. Generic Nature of the Condensed States of Proteins. Nat Cell Biol 2021, 23, 587–594. 10.1038/s41556-021-00697-8.

(3) Gabryelczyk, B.; Cai, H.; Shi, X.; Sun, Y.; Swinkels, P. J. M.; Salentinig, S.; Pervushin, K.; Miserez, A. Hydrogen Bond Guidance and Aromatic Stacking Drive Liquid-Liquid Phase Separation of Intrinsically Disordered Histidine-Rich Peptides. Nat Commun 2019, 10, 5465. 10.1038/s41467-019-13469-8.

(4) Malay, A.; Suzuki, T.; Katashima, T.; Kono, N.; Arakawa, K.; Numata, K. Spider Silk Self-Assembly via Modular Liquid-Liquid Phase Separation and Nanofibrillation. Sci Adv 2020, 6 (45), eabb6030.

(5) Tan, Y.; Hoon, S.; Guerette, P. A.; Wei, W.; Ghadban, A.; Hao, C.; Miserez, A.; Waite, J. H. Infiltration of Chitin by Protein Coacervates Defines the Squid Beak Mechanical Gradient. Nat Chem Biol 2015, 11, 488–495. 10.1038/nchembio.1833.

(6) Zbinden, A.; Pérez-Berlanga, M.; De Rossi, P.; Polymenidou, M. Phase Separation and Neurodegenerative Diseases: A Disturbance in the Force. Dev Cell 2020, 55, 45–68. 10.1016/j.devcel.2020.09.014.

(7) Shen, Y.; Ruggeri, F. S.; Vigolo, D.; Kamada, A.; Qamar, S.; Levin, A.; Iserman, C.; Alberti, S.; George-Hyslop, P. S.; Knowles, T. P. J. Biomolecular Condensates Undergo a Generic Shear-Mediated Liquid-to-Solid Transition. Nat Nanotechnol 2020, 15, 841–847. 10.1038/s41565-020-0731-4.

(8) Xu, Y.; Zhu, H.; Denduluri, A.; Ou, Y.; Erkamp, N. A.; Qi, R.; Shen, Y.; Knowles, T. P. J. Recent Advances in Microgels: From Biomolecules to Functionality. Small 2022, 18, e2200180. 10.1002/smll.202200180.

(9) Lipiński, W. P.; Visser, B. S.; Robu, I.; Fakhree, M. A. A.; Lindhoud, S.; Claessens, M. M. A. E.; Spruijt, E. Biomolecular Condensates Can Both Accelerate and Suppress Aggregation of α-Synuclein. Sci Adv 2022, 8, eabq6495. 10.1126/sciadv.abq6495.

(10) Gabryelczyk, B.; Philips, M.; Low, K.; Venkatraman, A.; Kannaian, B.; Alag, R.; Linder, M.; Pervushin, K.; Miserez, A. Liquid-Liquid Phase Separation Protects Amyloidogenic and Aggregation-Prone Peptides in Heterologous Expression Systems. Protein Science 2022, 31, e4292. 10.1101/2021.05.14.443401.

(11) Kanaan, N. M.; Hamel, C.; Grabinski, T.; Combs,B. Liquid-Liquid Phase Separation Induces Pathogenic Tau Conformations in Vitro. Nat Commun 2020, 11, 2809. 10.1038/s41467-020-16580-3.

(12) Ray, S.; Singh, N.; Kumar, R.; Patel, K.; Pandey, S.; Datta, D.; Mahato, J.; Panigrahi, R.; Navalkar, A.; Mehra, S.; Gadhe, L.; Chatterjee, D.; Sawner, A. S.; Maiti, S.; Bhatia, S.; Gerez, J. A.; Chowdhury, A.; Kumar, A.; Padinhateeri, R.; Riek, R.; Krishnamoorthy, G.; Maji, S. K. α-Synuclein Aggregation Nucleates through Liquid–Liquid Phase Separation. Nat Chem 2020, 12, 705–716. 10.1038/s41557-020-0465-9.

(13) Shen, Y.; Chen, A.; Wang, W.; Shen, Y.; Ruggeri, F. S.; Aime, S.; Wang, Z.; Qamar, S.; Espinosa, J. R.; Garaizar, A.; St George-Hyslop, P.; Collepardo-Guevara, R.; Weitz, D. A.; Vigolo, D.; Knowles, T. P. J. The Liquid-to-Solid Transition of FUS Is Promoted by the Condensate Surface. Proc Natl Acad Sci U S A 2023, 120, e2301366120. 10.1073/pnas.2301366120.

(14) Wei, S. P.; Qian, Z. G.; Hu, C. F.; Pan, F.; Chen, M. T.; Lee, S. Y.; Xia, X. X. Formation and Functionalization of Membraneless Compartments in Escherichia Coli. Nat Chem Biol 2020, 16, 1–6. 10.1038/s41589-020-0579-9.

(15) Vollrath, F.; Knight, D. P. Liquid Crystalline Spinning of Spider Silk. Nature 2001, 410 (6828), 541–548. 10.1038/35069000.

(16) Slotta, U. K.; Rammensee, S.; Gorb, S.; Scheibel, T. An Engineered Spider Silk Protein Forms Microspheres. Angewandte Chemie - International Edition 2008, 47, 4592–4594. 10.1002/anie.200800683.

(17) Mohammadi, P.; Christopher, J.; Beaune, G.; Engelhardt, P.; Kamada, A.; Timonen, J. V. I.; Knowles, T. P. J.; Penttila, M.; Linder, M. B. Controllable Coacervation of Recombinantly Produced Spider Silk Protein Using Kosmotropic Salts. J Colloid Interface Sci 2020, 560, 149–160. 10.1016/j.jcis.2019.10.058.

(18) Mohammadi, P.; Aranko, A. S.; Lemetti, L.; Cenev, Z.; Zhou, Q.; Virtanen, S.; Landowski, C. P.; Penttilä, M.; Fischer, W. J.; Wagermaier, W.; Linder, M. B. Phase Transitions as Intermediate Steps in the Formation of Molecularly Engineered Protein Fibers. Commun Biol 2018, 1, 86. 10.1038/s42003-018-0090-y.

(19) Breslauer, D. N.; Muller, S. J.; Lee, L. P. Generation of Monodisperse Silk Microspheres Prepared with Microfluidics. Biomacromolecules 2010, 11, 643–647. 10.1021/bm901209u.

(20) Toprakcioglu, Z.; Knowles, T. P. J. Shear-Mediated Sol-Gel Transition of Regenerated Silk Allows the Formation of Janus-like Microgels. Sci Rep 2021, 11, 6673. 10.1038/s41598-021-85199-1.

(21) Lammel, A.; Schwab, M.; Hofer, M.; Winter, G.; Scheibel, T. Recombinant Spider Silk Particles as Drug Delivery Vehicles. Biomaterials 2011, 32, 2233–2240. 10.1016/j.biomaterials.2010.11.060.

(22) Jawerth, L.; Fischer-Friedrich, E.; Saha, S.; Wang, J.; Franzmann, T.; Zhang, X.; Sachweh, J.; Ruer, M.; Ijavi, M.; Saha, S.; Mahamid, J.; Hyman, A. A.; Jülicher, F. Protein Condensates as Aging Maxwell Fluids. Science (1979) 2020, 370, 1317–1323. 10.1126/science.aaw4951.

(23) Wang, J.; Choi, J. M.; Holehouse, A. S.; Lee, H. O.; Zhang, X.; Jahnel, M.; Maharana, S.; Lemaitre, R.; Pozniakovsky, A.; Drechsel, D.; Poser, I.; Pappu, R. V.; Alberti, S.; Hyman, A. A. A Molecular Grammar Governing the Driving Forces for Phase Separation of Prion-like RNA Binding Proteins. Cell 2018, 174, 688–699. 10.1016/j.cell.2018.06.006.

(24) Chokkalingam, V.; Weidenhof, B.; Krämer, M.; Maier, W. F.; Herminghaus, S.; Seemann, R. Optimized Droplet-Based Microfluidics Scheme for Sol-Gel Reactions. Lab Chip 2010, 10, 1700–1705. 10.1039/b926976b.

(25) Xu, Y.; Qi, R.; Zhu, H.; Li, B.; Shen, Y.; Krainer, G.; Klenerman, D.; Knowles, T. P. J. Liquid–Liquid Phase-Separated Systems from Reversible Gel–Sol Transition of Protein Microgels. Advanced Materials 2021, 33, e2008670. 10.1002/adma.202008670.

(26) Shimanovich, U.; Song, Y.; Brujic, J.; Shum, H. C.; Knowles, T. P. J. Multiphase Protein Microgels. Macromol Biosci 2015, 15, 501–508. 10.1002/mabi.201400366.

(27) Schneider, N.; Wieland, F. G.; Kong, D.; Fischer, A. A. M.; Hörner, M.; Timmer, J.; Ye, H.; Weber, W. Liquid-Liquid Phase Separation of Light-Inducible Transcription Factors Increases Transcription Activation in Mammalian Cells and Mice. Sci Adv 2021, 7, eabd3568. 10.1126/sciadv.abd3568.

(28) Rising, A.; Johansson, J. Toward Spinning Artificial Spider Silk. Nature Chemical Biology. 2015. 10.1038/nchembio.1789.

(29) Andersson, M.; Jia, Q.; Abella, A.; Lee, X. Y.; Landreh, M.; Purhonen, P.; Hebert, H.; Tenje, M.; Robinson, C. V.; Meng, Q.; Plaza, G. R.; Johansson, J.; Rising, A. Bio-mimetic Spinning of Artificial Spider Silk from a Chimeric Minispidroin. Nat Chem Biol 2017, 13 (3), 262–264. 10.1038/nchembio.2269.

(30) Arndt, T.; Jaudzems, K.; Shilkova, O.; Francis, J.; Johansson, M.; Laity, P.; Sahin, C.; Chatterjee, U.; Kronqvist, N.; Barajas-Ledesma, E.; Kumar, R.; Chen, G.; Strömberg, R.; Abelein, A.; Langton, M.; Landreh, M.; Barth, A.; Holland, C.; Johansson, J.; Rising, A. Spidroin N-Terminal Domain Forms Amyloid-like Fibril Based Hydrogels and Provides a Protein Immobilization Platform. Nat Commun 2022, 13, 4695.

(31) Arndt, T.; Chatterjee, U.; Shilkova, O.; Francis, J.; Lundkvist, J.; Johansson, D.; Schmuck, B.; Greco, G.; Nordberg, Å. E.; Li, Y.; Wahlberg, L. U.; Langton, M.; Johansson, J.; Götherström, C.; Rising, A. Tuneable Recombinant Spider Silk Protein Hydrogels for Drug Release and 3D Cell Culture. Adv Funct Mater 2023. 10.1002/adfm.202303622.

(32) Leppert, A.; Chen, G.; Lama, D.; Sahin, C.; Railaite, V.; Shilkova, O.; Arndt, T.; Marklund, E. G.; Lane, D. P.; Rising, A.; Landreh, M. Liquid-Liquid Phase Separation Primes Spider Silk Proteins for Fiber Formation via a Conditional Sticker Domain. Nano Lett 2023, 23, 5836–5841.

(33) Greco, G.; Arndt, T.; Schmuck, B.; Francis, J.; Bäcklund, F. G.; Shilkova, O.; Barth, A.; Gonska, N.; Seisenbaeva, G.; Kessler, V.; Johansson, J.; Pugno, N. M.; Rising, A. Tyrosine Residues Mediate Supercontraction in Biomimetic Spider Silk. Commun Mater 2021, 2, 43. 10.1038/s43246-021-00147-w.

(34) Aggarwal, N.; Eliaz, D.; Cohen, H.; Rosenhek-Goldian, I.; Cohen, S. R.; Kozell, A.; Mason, T. O.; Shimanovich, U. Protein Nanofibril Design via Manipulation of Hydrogen Bonds. Commun Chem 2021, 4, 62. 10.1038/s42004-021-00494-2.

(35) Alberti, S.; Gladfelter, A.; Mittag, T. Considerations and Challenges in Studying Liquid-Liquid Phase Separation and Biomolecular Condensates. Cell 2019, 176, 419–434. 10.1016/j.cell.2018.12.035.

(36) Arndt, T.; Greco, G.; Schmuck, B.; Bunz, J.; Shilkova, O.; Francis, J.; Pugno, N. M.; Jaudzems, K.; Barth, A.; Johansson, J.; Rising, A. Engineered Spider Silk Proteins for Biomimetic Spinning of Fibers with Toughness Equal to Dragline Silks. Adv Funct Mater 2022, 32, 2200986.

(37) Hammarström, P.; Simon, R.; Nyström, S.; Konradsson, P.; Åslund, A.; Nilsson, K. P. R. A Fluorescent Pentameric Thiophene Derivative Detects in Vitro-Formed Pre-fibrillar Protein Aggregates. Biochemistry 2010, 49, 6838–6845. 10.1021/bi100922r.

(38) Klingstedt, T.; Shirani, H.; Åslund, K. O. A.; Cairns, N. J.; Sigurdson, C. J.; Goedert, M.; Nilsson, K. P. R. The Structural Basis for Optimal Performance of Oligothio-phene-Based Fluorescent Amyloid Ligands: Conformational Flexibility Is Essential for Spectral Assignment of a Diversity of Protein Aggregates. Chemistry - A European Journal 2013, 19, 10179–10192. 10.1002/chem.201301463.

(39) Garaizar, A.; Espinosa, J. R.; Joseph, J. A.; Krainer, G.; Shen, Y.; Knowles, T. P. J.; Collepardo-Guevara, R. Aging Can Transform Single-Component Protein Condensates into Multiphase Architectures. Proc Natl Acad Sci U S A 2022, 119, e2119800119. 10.1073/pnas.2119800119.

(40) Blazquez, S.; Sanchez-Burgos, I.; Ramirez, J.; Higginbotham, T.; Conde, M. M.; Collepardo-Guevara, R.; Tejedor, A. R.; Espinosa, J. R. Location and Concentration of Aromatic-Rich Segments Dictates the Percolating Inter-Molecular Network and Viscoelastic Properties of Ageing Condensates. Advanced Science 2023, 10, e2207742. 10.1002/advs.202207742.

(41) Andersson, M.; Chen, G.; Otikovs, M.; Landreh, M.; Nordling, K.; Kronqvist, N.; Westermark, P.; Jörnvall, H.; Knight, S.; Ridderstråle, Y.; Holm, L.; Meng, Q.; Jaudzems, K.; Chesler, M.; Johansson, J.; Rising, A. Carbonic Anhydrase Generates CO2 and H+ That Drive Spider Silk Formation Via Opposite Effects on the Terminal Domains. PLoS Biol 2014, 12 (8), e1001921. 10.1371/journal.pbio.1001921.

(42) Kaldmäe, M.; Leppert, A.; Chen, G.; Sarr, M.; Sahin, C.; Nordling, K.; Kronqvist, N.; Gonzalvo-Ulla, M.; Fritz, N.; Abelein, A.; Laín, S.; Biverstål, H.; Jörnvall, H.; Lane, D. P.; Rising, A.; Johansson, J.; Landreh, M. High Intracellular Stability of the Spidroin N-Terminal Domain in Spite of Abundant Amyloidogenic Segments Revealed by in-Cell Hydrogen/Deuterium Exchange Mass Spectrometry. FEBS Journal 2020, 287 (13), 2823–2833.

(43) Kenworthy, A. K. What’s Past Is Prologue: FRAP Keeps Delivering 50 Years Later. Biophysical Journal 2023, 122, 3577–3586. 10.1016/j.bpj.2023.05.016.

(44) Taylor, N. O.; Wei, M. T.; Stone, H. A.; Brangwynne, C. P. Quantifying Dynamics in Phase-Separated Condensates Using Fluorescence Recovery after Photobleaching. Biophysical Journal 2019, 117, 1285–1300. 10.1016/j.bpj.2019.08.030.

(45) Liang, C. Q.; Wang, L.; Luo, Y. Y.; Li, Q. Q.; Li, Y. M. Capturing Protein Droplets: Label-Free Visualization and Detection of Protein Liquid-Liquid Phase Separation with an Aggregation-Induced Emission Fluorogen. Chemical Communications 2021, 57, 3805–3808. 10.1039/d1cc00947h.

(46) Pansieri, J.; Josserand, V.; Lee, S. J.; Rongier, A.; Imbert, D.; Sallanon, M. M.; Kövari, E.; Dane, T. G.; Vendrely, C.; Chaix-Pluchery, O.; Guidetti, M.; Vollaire, J.; Fertin, A.; Usson, Y.; Rannou, P.; Coll, J. L.; Marquette, C.; Forge, V. Ultraviolet–Visible–near-Infrared Optical Properties of Amyloid Fibrils Shed Light on Amyloidogenesis. Nature Photonics 2019, 13, 473–479. 10.1038/s41566-019-0422-6.

(47) Monti, A.; Bruckmann, C.; Blasi, F.; Ruvo, M.; Vitagliano, L.; Doti, N. Amyloid-like Prep1 Peptides Exhibit Reversible Blue-Green-Red Fluorescencein Vitroand in Living Cells. Chemical Communications 2021, 57, 3720–3723. 10.1039/d1cc01145f.

(48) Pinotsi, D.; Grisanti, L.; Mahou, P.; Gebauer, R.; Kaminski, C. F.; Hassanali, A.; Kaminski Schierle, G. S. Proton Transfer and Structure-Specific Fluorescence in Hydrogen Bond-Rich Protein Structures. Journal of the American Chemical Society 2016, 138, 3046–3057. 10.1021/jacs.5b11012.

(49) Shimanovich, U.; Pinotsi, D.; Shimanovich, K.; Yu, N.; Bolisetty, S.; Adamcik, J.; Mezzenga, R.; Charmet, J.; Vollrath, F.; Gazit, E.; Dobson, C. M.; Schierle, G. K.; Holland, C.; Kaminski, C. F.; Knowles, T. P. J. Biophotonics of Native Silk Fibrils. Macromolecular Bioscience 2018, 18, e1700295. 10.1002/mabi.201700295.

(50) Apter, B.; Lapshina, N.; Barhom, H.; Fainberg, B.; Handelman, A.; Accardo, A.; Diaferia, C.; Ginzburg, P.; Morelli, G.; Rosenman, G. Fluorescence Phenomena in Amyloid and Amyloidogenic Bionanostructures. Crystals 2020, 10, 668. 10.3390/cryst10080668.

(51) Del Mercato, L. L.; Pompa, P. P.; Maruccio, G.; Della Torre, A.; Sabella, S.; Tamburro, A. M.; Cingolani, R.; Rinaldi, R. Charge Transport and Intrinsic Fluorescence in Amyloid-like Fibrils. Proceedings of the National Academy of Sciences of the United States of America 2007, 104, 18019–18024. 10.1073/pnas.0702843104.

(52) Pinotsi, D.; Buell, A. K.; Dobson, C. M.; Kaminski Schierle, G. S.; Kaminski, C. F. A Label-Free, Quantitative Assay of Amyloid Fibril Growth Based on Intrinsic Fluorescence. ChemBioChem 2013, 14, 846–850. 10.1002/cbic.201300103.

(53) Klein, I. A.; Boija, A.; Afeyan, L. K.; Hawken, S. W.; Fan, M.; Dall’Agnese, A.; Oksuz, O.; Henninger, J. E.; Shrinivas, K.; Sabari, B. R.; Sagi, I.; Clark, V. E.; Platt, J. M.; Kar, M.; McCall, P. M.; Zamudio, A. V.; Manteiga, J. C.; Coffey, E. L.; Li, C. H.; Hannett, N. M.; Guo, Y. E.; Decker, T. M.; Lee, T. I.; Zhang, T.; Weng, J. K.; Taatjes, D. J.; Chakraborty, A.; Sharp, P. A.; Chang, Y. T.; Hyman, A. A.; Gray, N. S.; Young, R. A. Partitioning of Cancer Therapeutics in Nuclear Condensates. Science (1979) 2020, 368, 1386–1392. 10.1126/science.aaz4427.

(54) Stengel, D.; Saric, M.; Johnson, H. R.; Schiller, T.; Diehl, J.; Chalek, K.; Onofrei, D.; Scheibel, T.; Holland, G. P. Tyrosine’s Unique Role in the Hierarchical Assembly of Recombinant Spider Silk Proteins: From Spinning Dope to Fibers. Biomacromolecules 2022, 24, 1463–1474. 10.1021/acs.biomac.2c01467.

(55) Gabryelczyk, B.; Sammalisto, F. E.; Gandier, J. A.; Feng, J.; Beaune, G.; Timonen, J. V. I.; Linder, M. B. Recombinant Protein Condensation inside E. Coli Enables the Development of Building Blocks for Bioinspired Materials Engineering – Biomimetic Spider Silk Protein as a Case Study. Mater Today Bio 2022, 17, 100492. 10.1016/j.mtbio.2022.100492.

(56) Avni, A.; Joshi, A.; Walimbe, A.; Pattanashetty, S. G.; Mukhopadhyay, S. Single-Droplet Surface-Enhanced Raman Scattering Decodes the Molecular Determinants of Liquid-Liquid Phase Separation. Nature Communications 2022, 13, 4378. 10.1038/s41467-022-32143-0.

