## Supplementary material for "Controlling Drug Partitioning in Individual Protein Condensates through Laser-Induced Microscale Phase Transitions": FIgure S1

| | |
|---|---|
| Experimental procedures | 2 - 3 |
| Supporting Figures 1-4 | 3 - 7 |
| Supporting References | 8 |

**Experimental Procedures**

**Chemicals**
All reagents were purchased from Sigma-Aldrich if not stated otherwise.

**Sequences**

| | |
|---|---|
| NT2RepCT | MGHHHHHHMSHTTPWTNPGLAENFMNSFMQGLSSMPGFTASQLDDMSTIAQSMVQSIQSL<br>AAQGRTSPNKLQALNMAFASSMAEIAASEEGGGSLSTKTSSIASAMSNAFLQTTGVVNQPFI<br>NEITQLVSMFAQAGMNDVSAGNSGRGQGG**Y**GQGSGGNAAAAAAAAAAAAAAAGQGGQG<br>G**Y**GRQSQGAGSAAAAAAAAAAAAAAGSGQGG**Y**GGQGQGG**Y**GQSGSVTSGGYGYGTSAA<br>AGAGVAAGSYAGAVNRLSSAEAASRVSSNIAAIASGGASALPSVISNIYSGVVASGVSSNEA<br>LIQALLELLSALVHVLSSASIGNVSSVGVDSTLNVVQDSVGQYVG |
| NT2RepCT$^{YF}$ | MGHHHHHHMSHTTPWTNPGLAENFMNSFMQGLSSMPGFTASQLDDMSTIAQSMVQSIQSL<br>AAQGRTSPNKLQALNMAFASSMAEIAASEEGGGSLSTKTSSIASAMSNAFLQTTGVVNQPFI<br>NEITQLVSMFAQAGMNDVSAGNSGRGQGG**F**GQGSGGNAAAAAAAAAAAAAAAGQGGQG<br>G**F**GRQSQGAGSAAAAAAAAAAAAAAGSGQGG**F**GGQGQGG**F**GQSGSVTSGGYGYGTSAA<br>AGAGVAAGSYAGAVNRLSSAEAASRVSSNIAAIASGGASALPSVISNIYSGVVASGVSSNEA<br>LIQALLELLSALVHVLSSASIGNVSSVGVDSTLNVVQDSVGQYVG |

**Protein expression and purification**
NT2RepCT and NT2RepCT$^{YF}$ was expressed and purified as previously described except that sonication (5 min, 2 sec on/ 2 sec off, 30% amplitude) instead of a high-pressure cell disrupter was used for protein solubilization.[51] Purified protein was dialyzed against deionized water and stored at -80°C. $^{15}$N-labeled NT2RepCT$^{YF}$ was expressed in *E. coli* BL21 (DE3) cells using 2x M9 medium. 20 ml start culture containing 2x M9 medium and 75 µg/ml kanamycin was inoculated with NT2RepCT$^{YF}$ from a previously prepared glycerol stock stored in Luria broth (LB) medium and incubated overnight at 37°C. Cells from the start culture (5% volume of the main culture) were transferred in fresh 2x M9 medium containing 75 µg/ml kanamycin and incubated at 31°C until the OD600 nm reached 0.9. For protein expression the incubation temperature was lowered to 20°C, Isopropyl b-D-1-thiogalactopyranoside (IPTG) was added to 0.5 mM, and cells were incubated overnight. Protein purification was carried out as for unlabeled NT2RepCT$^{YF}$. hTau was expressed and purified as previously described and stored at -20°C in 20 mM sodium phosphate buffer pH 7.4 with 0.2 mM EDTA.[52] Met-Ab42 was expressed and purified as previously described.[53]

**Labeling**
Prior to the labeling reaction protein stocks were buffer exchanged to 10 mM sodium phosphate pH 8.3 using slide-A-Lyzer mini dialysis tubes (Thermo Scientific, 88401). Freeze dried Atto655 NHS-ester dye (Sigma-Aldrich, 76245) was dissolved in a small volume of acetonitrile, mixed in a dye to protein ratio of 2:1 mole/mole and incubated at room temperature for 30 min. Unbound dye was removed using Micro Bio-Spin 6 columns (Bio-Rad) against deionized water. The absorbance at 280 nm and 663 nm were measured and used to calculate the degree of labeling (DOL).

**Droplet formation and maturation**
Protein droplets of NT2RepCT$^{YF}$ were formed by mixing 25 µM protein in 0.5 M potassium phosphate pH 8 in low binding tubes (Axygen) if not stated otherwise. Samples have been dispensed into black 96-well low-binding polystyrene microplates with a transparent bottom (Corning). Samples have been analyzed directly after 30 min sedimentation time and after maturation at 37°C for 24 h or at room temperature for at least 72 h if not stated otherwise. For maturation experiments plates have been sealed with cover film to avoid evaporation. Droplet formation of hTau was induced without co-factors (such as crowding agents or RNA) by buffer exchanging 122 µM hTau (stored in 20 mM sodium phosphate, pH 7.4 with 0.2 mM EDTA) to 25 mM ammonium acetate pH 8 and further dilution to 10 µM in deionized water.

**Fluorescence emission spectroscopy and kinetics**
Thioflavin T (ThT) and pentameric formyl thiophene acetic acid (pFTAA) fluorescence emission was recorded using a SPARK 20M plate reader (Tecan). ThT kinetics were monitored using an excitation wavelength of 448 nm, an emission wavelength of 485 nm, 5 nm bandwidths and a gain of 150. Samples containing 25 µM NT2RepCT$^{YF}$ were prepared with 10 µM ThT in either deionized water or 0.5 M potassium phosphate pH 8.0 buffer. Five replicates, each containing 40 µl of reactant volume, were dispensed into black 386-well low-binding polystyrene microplates with a transparent bottom (Corning) and sealed with

transparent cover film to avoid evaporation during incubation at 37°C. To measure pFTAA fluorescence protein samples were supplemented with 0.4 µM pFTAA prior to maturation in either deionized water or 0.5 M potassium phosphate pH 8.0. Emission spectra between 470 nm and 670 nm were recorded in 1 nm increments at 25 °C using an excitation wavelength of 455 nm, 7.5 nm bandwidths and a gain of 170. As amyloid control, amyloid-b fibrils were generated from a 3 µM monomeric amyloid-b solution in 20 mM sodium phosphate pH 8 with 0.2 mM EDTA.

**Fluorescence microscopy and FRAP**

For fluorescence microscopy and FRAP experiments samples have been supplemented with either 10 µM ThT, 25 µM DroProbe, or 25 µM Mitoxantrone prior to droplet formation. Fluorescence microscopy images in Figure 1 were acquired using a Nikon Eclipse Ti series inverted microscope (Nikon) equipped with a Crest X-light V2 series confocal unit (Nikon) using a Plan Apo 40x objective (Nikon) and a Zyla sCMOS camera (Andor). ThT incubated samples were excited with a 470 nm laser. To compare different time points, laser power and exposure were kept constant throughout each experiment.

FRAP experiments were performed on a LSM980-Airy microscope (Zeiss) equipped with an Airy detector 2 using a C-Apochromat 40X water objective. Fluorescence was excited using a 639 nm (Atto655 and Mitoxantrone) and 405 nm (ThT and DroProbe) wavelengths, a pinhole of 49 µm, a master gain of 700 V, at low laser power (3 – 5%) and short exposure time. Photobleaching was achieved in a selected region of interest (~ 1 µm diameter) by increasing the laser power to 100% for 50 iterations and lowering the scan speed from 7 to 5 during bleaching in a selected region of interest. Fluorescence recovery was recorded using the same settings as before bleaching. Autofluorescence of fresh and matured NT2RepCT$^{YF}$ droplets, and hTau has been performed using the same microscope and objective that has been used for FRAP experiments. Droplets were excited using 405 nm (30% laser power), 561 nm (50% laser power) or 639 nm (100% laser power) wavelengths and the emission range was set to 431 nm – 560 nm, 572 nm – 736 nm and 644 nm – 739 nm, respectively. The pinhole was set to 49 µm, the master gain to 850 V and the digital gain to 1. To compare different samples all settings were kept constant throughout each experiment, only the laser power has been adjusted dependent on the laser line. Images were processed using ImageJ.

**Live cell imaging**

*E.coli* cell cultures suspended in growth media were placed on a microscope coverslip coated with poly-D-Lysine (A3890401 Thermo Fisher Scientific). Images were acquired using a Nikon Ti-E inverted microscope equipped with a 60×/1.4 objective lens, and Crest OpticsX-lightV3 spinning disk confocal head (operated in the widefield mode). The fluorescence signal of eGFP-NT2RepCT was excited by continuous illumination on samples using 470 nm light from an LDI Laser Diode Illuminator (89 North) at 1% power level with an exposure time of 50 ms, while emission light between 485 nm and 535 nm was collected. A Gataca Systems iLas 2 unit coupled to a 100 mW OBIS LX 405 nm laser was used to generate a circular spot with a diameter of approximately 1 µm for photobleaching. The photobleaching on samples was carried out at a fixed 4% laser power level and at a controlled exposure time of 0.1 s. After photobleaching, the fluorescence images of samples were acquired using the above setting but with an exposure time of 50 ms to minimize imaging-induced photobleaching. The same setting of FRAP was used for E.coli cells grown at 18°C and 37°C. Processing of the images and quantification of the fluorescence recovery were carried out using the Fiji (2.3.0) software with a plugin developed by Jay Unruh at the Stowers Institute for Medical Research (KansasCity, MO). [54]

**Sample preparation for native mass spectrometry**

To study the exchange of molecules between the soluble and dense phase of NT2RepCT$^{YF}$ by native MS, the following samples have been prepared. A fully soluble protein sample was prepared by mixing 100 µM NT2RepCT$^{YF}$ and 10 µM $^{15}$N-labeled NT2RepCT$^{YF}$ in 100 mM ammonium acetate pH 8. This sample was directly analyzed by nMS. Phase separation of 100 µM NT2RepCT$^{YF}$ was induced in 0.35 M potassium phosphate buffer pH 8 in low binding tubes (Axygen) and 50 µl dispensed in a low-bind 96-well plate. Subsequently, $^{15}$N -labeled NT2RepCT$^{YF}$ was added to a final concentration of 10 µM directly to the supernatant of phase separated NT2RepCT$^{YF}$ (fresh) or after NT2RepCT$^{YF}$ condensates have been aged for 72 h at RT (gelated). Immediately after addition of $^{15}$N-labeled NT2RepCT$^{YF}$ samples have been collected in low-bind tubes and centrifuged at 15000xg for 2 min at 25°C. The supernatants of fresh and gelated samples were collected, and buffer exchanged to 100 mM ammonium acetate pH 8 using Zeba Spin desalting columns, 7k MWCO (Thermo Fisher Scientific) and diluted 1:1 with 100 mM ammonium acetate pH 8 prior to MS analysis. The pellet from **freshly** formed condensates has been resuspended in 100 mM ammonium acetate and 5% acetonitrile added prior to MS analysis. The pellet sample of **matured** condensates was not amendable to MS analysis.

**Native Mass Spectrometry**

Native mass spectra were acquired on a Waters Synapt G1 travelling wave ion mobility mass spectrometer (Waters, UK) equipped with an offline nanospray source. The capillary voltage was set to 1.5 kV, the source pressure was maintained at 7.5 mbar, the sampling cone voltage set to 20 V, the source temperature was 30 °C, and the collision energy in the ion trap was 30 V. Spectra were visualized using MassLynx 4.1 (Waters, UK).

**Supporting Figures**

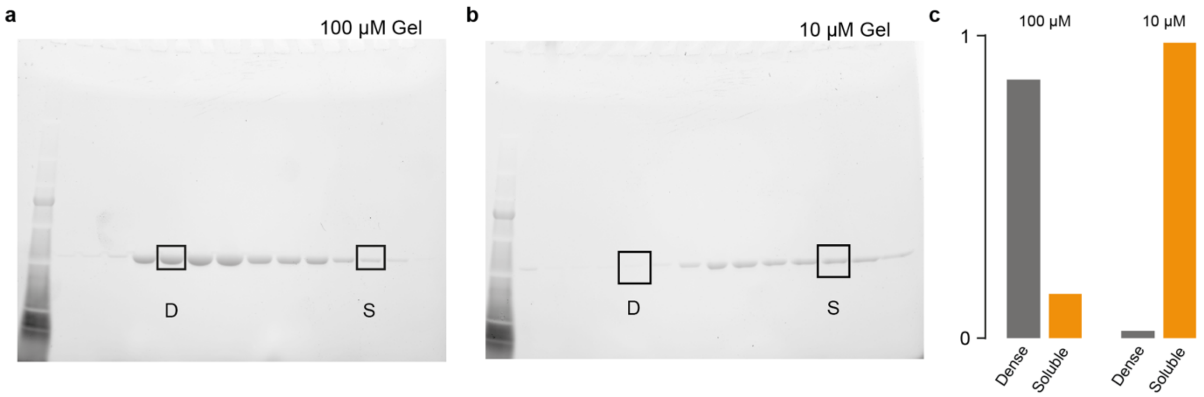


**Figure S1.** (a) and (b) Solubility of NT2RepCT$^{YF}$. The protein was incubated in 0.35 M $KPO_4$, pH 8 at concentrations of 100 μM or 10 μM. The dense (D) and soluble (S) fractions were separated by centrifugation and analyzed by SDS-PAGE. (c) Quantification of the band intensities (shown in rectangles in a and b) from the SDS-PAGE. At 100 μM, approximately 90% of the protein was in the dense phase, while at 10 μM, > 95% remained in the soluble phase.

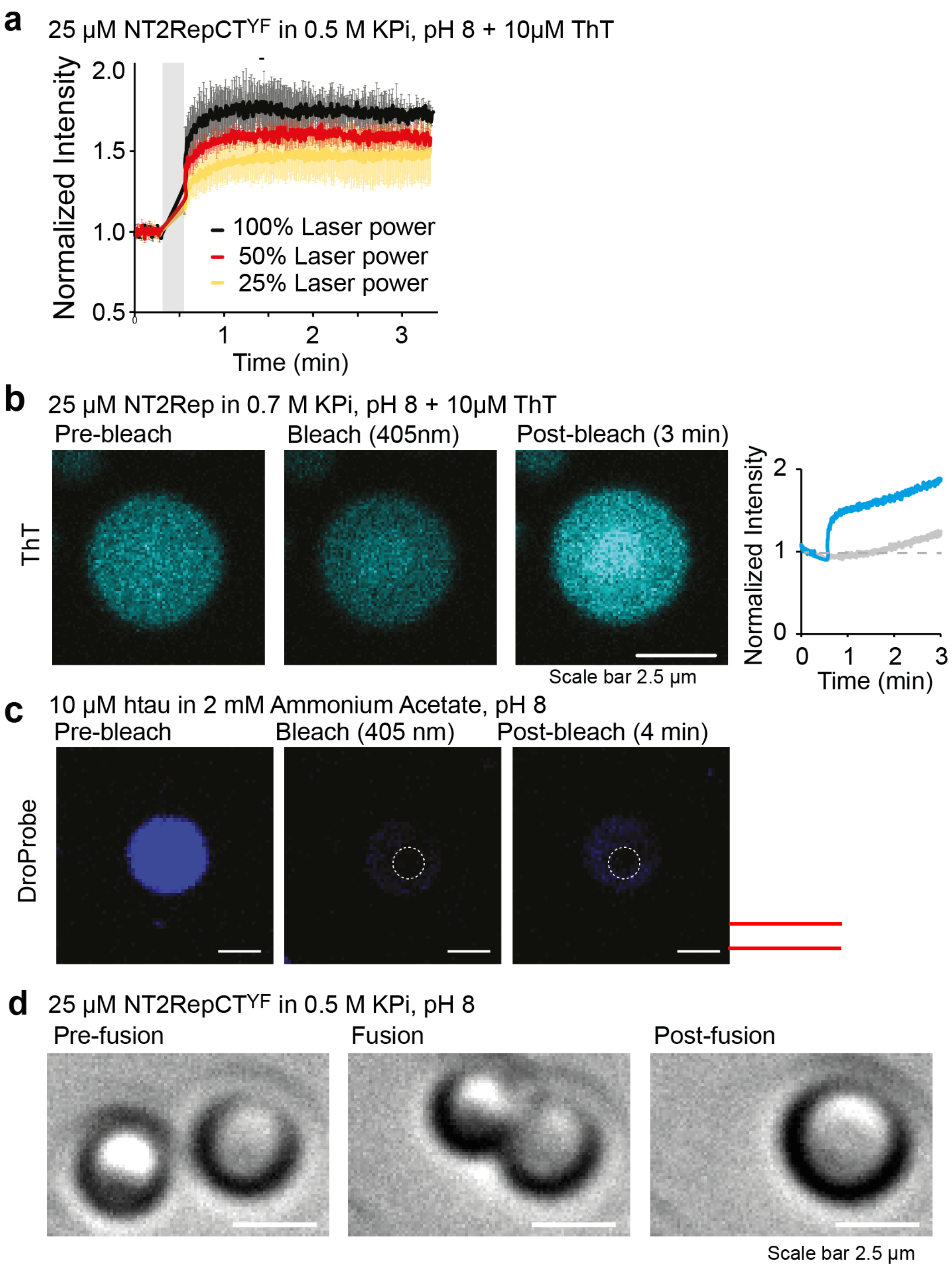


**Figure S2.** (a) ThT fluorescence traces for individual fresh droplets (25 µM) after photobleaching at a wavelength of 405 nm with laser intensities set to 25, 50, and 100%. Error bars indicate the standard deviation of three independent repeats. The bleaching regime is indicated by a grey bar. (b) Droplet formation by a truncated spidroin. 25 µM NT2Rep (without CTD) forms droplets in 0.7 M $KPO_4$, pH8. NT2Rep droplets display the same overshoot in ThT fluorescence after photobleaching as NT2RepCT$^{YF}$, indicating that gelation is driven by thermal denaturation of the NTD. Scale bar is 2.5 µm. Fluorescence traces (right) are shown for a bleached droplet (blue trace) and a neighboring, unbleached droplet (gray trace). (c) Photobleaching of htau droplets stained with DroProbe reagent using the same maximum intensity laser pulse settings as for gelation of NT2RepCT$^{YF}$ show complete bleaching and only minor fluorescence recovery, with no intensity overshoot. Scale bar is 5 µm. (d) NT2RepCT$^{YF}$ condensates are liquid prior to photobleaching, as judged by droplet fusion observed by brightfield microscopy. Scale bar is 2.5 µm.

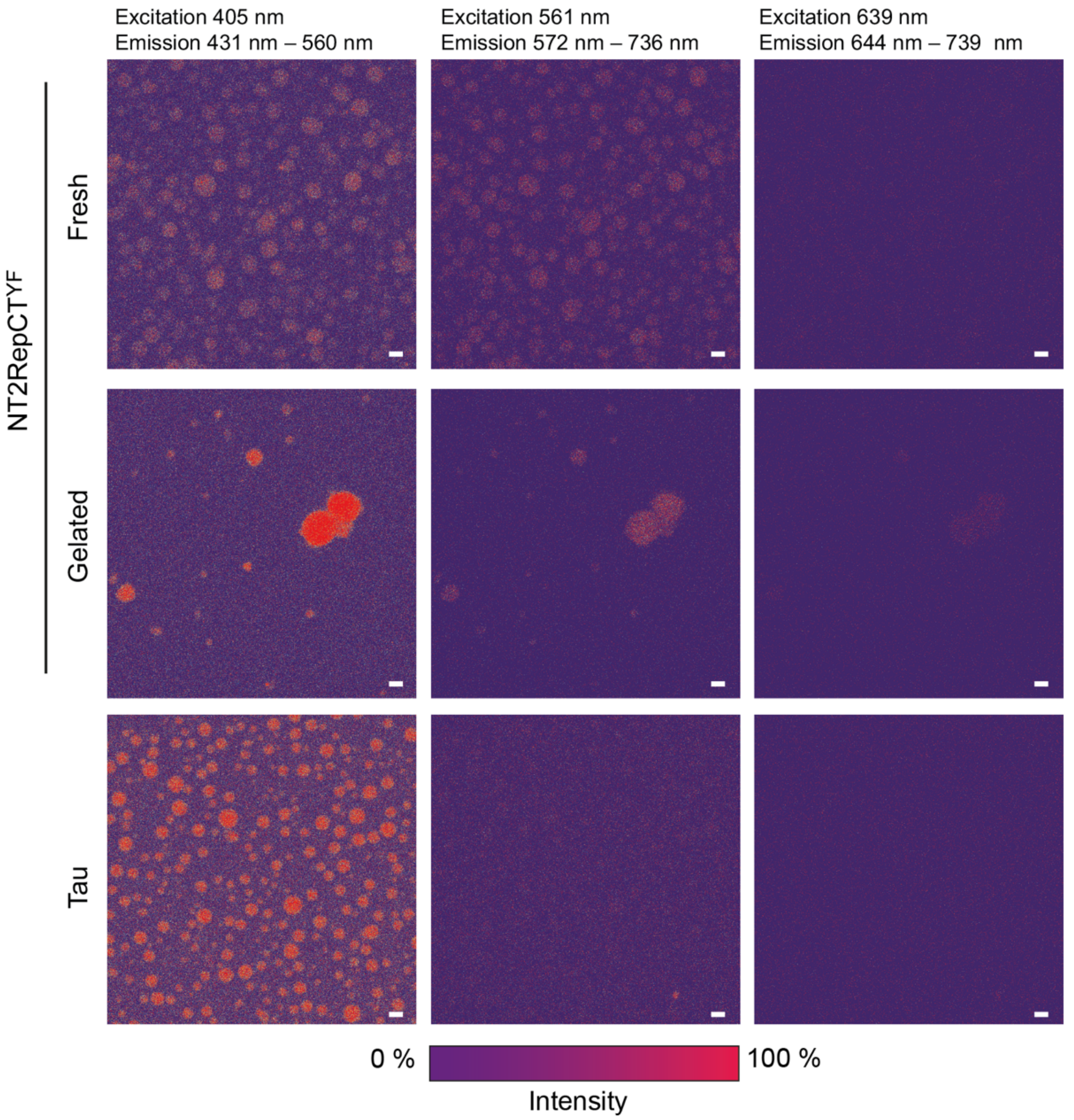


**Figure S3.** Confocal microscopy of fresh (top row) or gelated NT2RepCT$^{YF}$ droplets (middle row) shows red-shifted fluorescence upon excitation at λ 405 and λ561 nm, (emission λ 431 nm - 560 nm and λ 572 nm - 736 nm, respectively) which is significantly less pronounced upon excitation at λ 639 nm (emission λ 644 nm - 739 nm). Droplets of htau (bottom row) exhibit prominent fluorescence at λ 405, but not at λ 561 nm or λ 639 nm. Scale bars are 5 µm. All images are shown with the same color intensity scale below.

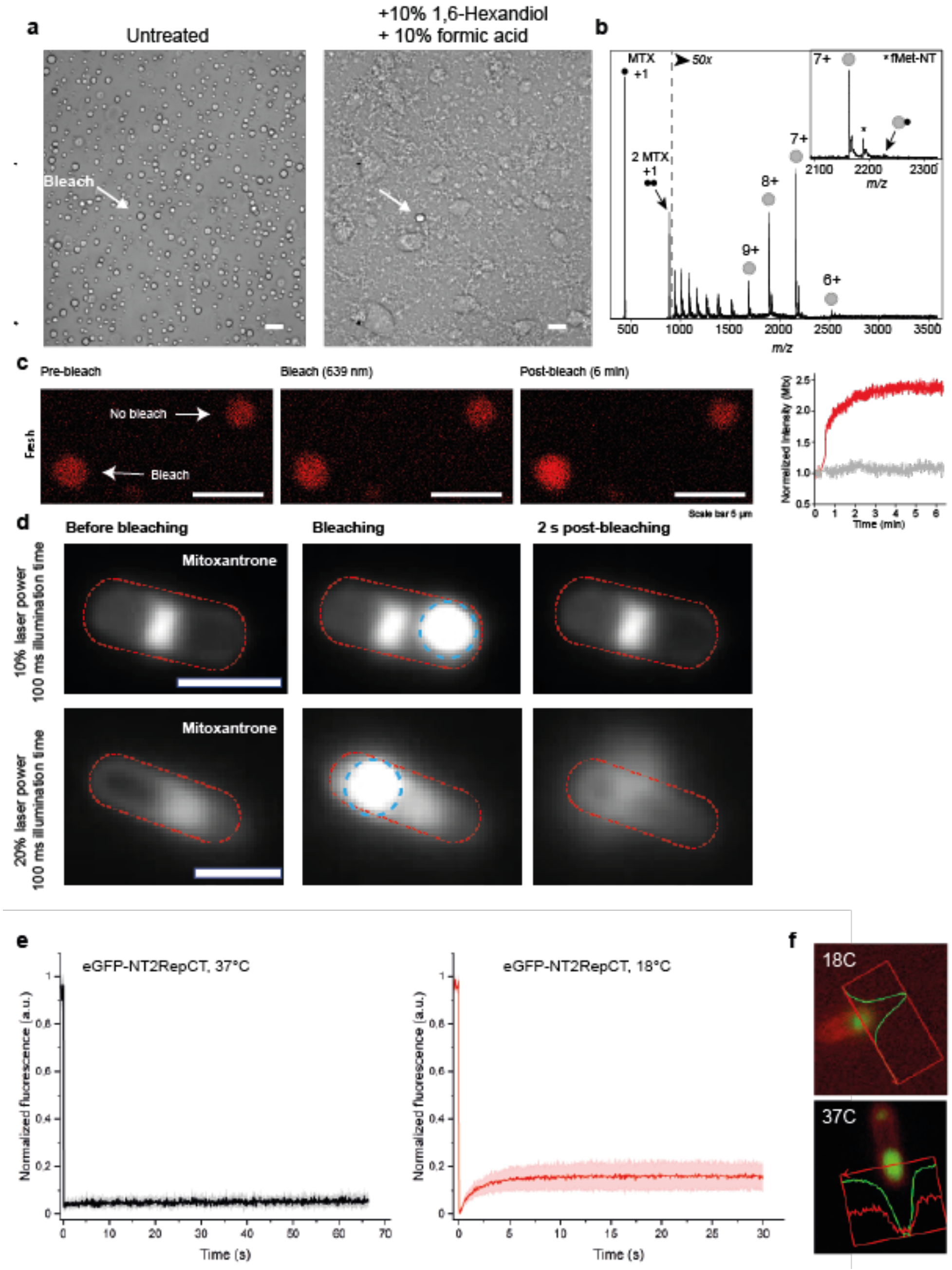


**Figure S4.** (a) Photobleaching of an individual droplet (arrow) induces resistance to 1,6-hexanediol and formic acid. Brightfield microscopy before (left) and after (right) addition of 10% 1,6-hexanediol and 10% formic acid are shown. Scale bars are 5 µm. (c) Photobleaching of wild-type NT2RepCT in the presence of Mitoxantrone shows post-bleach fluorescence overshoot as detected for $NT2RepCT^{YF}$. Scale bar is 5 µm. (d) Fluorescence microscopy images show the effects of increased laser power on the integrity of *E. coli*, indicating significant heating from high-energy FRAP. Top row: *E. coli* with intracellular mitoxantrone (white) before, during, and 2 s after a 100 ms laser pulse with 10% power shows no effect on cell integrity. Bottom row: Increasing the laser power to 20% results in cell lysis. Scale bars are 2 µM. (e) Intracellular eGFP-NT2RepCT condensates formed at 37°C show no recovery after photobleaching (left), whereas those formed at 18°C do, indicating liquid-like properties. The standard deviations of each condition are plotted in the form of a shaded area. At least four independent repeats were performed for both expression temperatures. (f) Images of the eGFP-NT2RepCT condensates used to quantify Mitoxantrone colocalization in Figure 3e in the main manuscript.

**References**


(1) Arndt, T.; Greco, G.; Schmuck, B.; Bunz, J.; Shilkova, O.; Francis, J.; Pugno, N. M.; Jaudzems, K.; Barth, A.; Johansson, J.; Rising, A. Engineered Spider Silk Proteins for Biomimetic Spinning of Fibers with Toughness Equal to Dragline Silks. *Adv. Funct. Mater.* **2022**, *32*, 2200986.

(2) Danis, C.; Despres, C.; Bessa, L. M.; Malki, I.; Merzougui, H.; Huvent, I.; Qi, H.; Lippens, G.; Cantrelle, F. X.; Schneider, R.; Hanoulle, X.; Smet-Nocca, C.; Landrieu, I. Nuclear Magnetic Resonance Spectroscopy for the Identification of Multiple Phosphorylations of Intrinsically Disordered Proteins. *J. Vis. Exp.* **2016**, *118*, 55001. https://doi.org/10.3791/55001.

(3) Leppert, A.; Tiiman, A.; Kronqvist, N.; Landreh, M.; Abelein, A.; Vukojević, V.; Johansson, J. Smallest Secondary Nucleation Competent Aβ Aggregates Probed by an ATP-Independent Molecular Chaperone Domain. *Biochemistry* **2021**, *60*, 678–688. https://doi.org/10.1021/acs.biochem.1c00003.

(4) Song, J.; Mizrak, A.; Lee, C. W.; Cicconet, M.; Lai, Z. W.; Tang, W. C.; Lu, C. H.; Mohr, S. E.; Farese, R. V.; Walther, T. C. Identification of Two Pathways Mediating Protein Targeting from ER to Lipid Droplets. *Nat. Cell Biol.* **2022**, *24*, 1364–1377. https://doi.org/10.1038/s41556-022-00974-0.
